# Spatio-temporal regulation of Dachsous proteins during *Drosophila* development

**DOI:** 10.1101/325977

**Authors:** Eva Revilla-Yates, Javier Sierra, Isabel Rodriguez

## Abstract

Transcriptional regulation is one of the main mechanisms involved in tissue morphogenesis to give rise to functional organs with characteristic shapes. The *Drosophila dachsous* (*ds*) gene plays a key role in tissue development and tumorigenesis by controlling planar cell polarity (PCP), tissue growth, patterning and mitochondrial activity, among other processes. Disturbance of *ds* expression during *Drosophila* development results in alterations of the function and morphology of a wide range of embryonic and larval tissues. Similarly, in humans, mutations in the *DCHS1* gene cause severe congenital malformations due to a global impairment affecting the normal formation of many tissues and organs. However, the transcriptional mechanism governing the expression of *ds* gene remains poorly understood. Here, we perform transcriptional analysis of *ds* expression and identify novel embryonic Ds proteins not expressed in larvae. The comparative analysis of Ds proteins and the exon expression pattern in/of two regulatory alleles such as *ds^D36^* and *ds^38K^* further suggests the existence of specific transcriptional *ds* variants at different stages. Furthermore, a search for regulatory elements that control the spatial and temporal pattern of *ds* revealed the presence of cis-regulatory elements located in the intronic regions, which regulate the expression of these Ds proteins. Finally, using the *Drosophila* wing as model to perform a functional analysis, we show that wing growth and PCP are differentially regulated by Ds proteins expressed in different regions of wing disc. The present findings reveal that the complex regulation of the *ds* gene ensures the expression of specific Ds protein isoforms at different developmental stages in order to activate the cell-specific molecular programs required for tissue morphogenesis.

## INTRODUCTION

Transcriptional regulation is one of the main mechanisms involved in tissue morphogenesis. Every aspect of the cellular function depends on the gene products expressed in the cell. The identification of the cis-acting elements and trans-acting factors that regulate gene expression in time and space is crucial to understand how specific cellular processes are coordinated to give rise to the variety of shapes specified during development. The *Drosophila dachsous* (*ds*) gene is an excellent candidate to gain knowledge into how the transcriptional regulation of a gene is able to coordinate a network of subordinate signalling pathways and transcription factors to guide different cellular processes within a cell population, including proliferation, oriented cell division, cell-cell communication and cell fate specification (reviewed in (Matis and Axelrod, 2013). During *Drosophila* development, *ds* is expressed in multiple cell types and its loss of function results in early lethality and abnormal size and shape of different embryonic and larval tissues such as the tracheal system (Chung et al., 2009), gut [Gonzalez-Morales et al. 2015], peripheral and central nervous system (Dearborn and Kunes, 2004); (Fabre et al., 2008) and imaginal discs (Rawls et al., 2002);(Rodriguez, 2004);(Bando et al., 2009). Furthermore, mutations in the *DCHS1* gene, the human homologue of *ds*, cause severe congenital malformations due to a global impairment in development affecting the normal formation of many tissues and organs including craniofacial, skeletal and limb malformations, hearing loss and heart valve and brain abnormalities suggesting that the altered expression of *DCHS1* by diverse polymorphisms can affect in a tissue-specific and developmentally specific manner (Cappello et al., 2013);(Sotos et al., 2017);(Clemenceau et al., 2017);(Zwaveling-Soonawala et al., 2018). The *ds* gene was initially identified as a member of the cadherin superfamily that controls the establishment of Planar Cell Polarity (PCP), a general mechanism required for tissue organization and organ formation (Clark et al., 1995). Subsequent genetic studies have shown the role of *ds* in patterning, tissue growth and mitochondrial activity regulating the activity of Hippo and JNK pathways and the expression of *dally* and *dally-like* (*dlp*) genes (Matakatsu and Blair, 2006); (Willecke et al., 2008). (Baena-Lopez et al., 2008);(Degoutin et al., 2013);(Revilla-Yates et al., 2015). Recently, we have reported the characterization of three novel Ds transcriptional isoforms expressed in the imaginal discs and brain of *Drosophila* larvae, in addition to the large transmembrane protein DsFL previously reported (Clark et al., 1995);(Revilla-Yates et al., 2015). These Ds proteins differ in their structural features, and whereas the DsEx and Ds1 isoforms are formed by 17 and 4 cadherin domains, respectively, the third isoform, DsIntra, is a small cytoplasmic protein that plays an essential role in mitochondrial activity (Revilla-Yates et al., 2015). However, it is unclear how the expression pattern of these Ds isoforms is regulated in the larval tissues, as well as their contribution to the control of patterning, growth and PCP. Key to understand how *ds* can control different cell functions over different windows of time during development is to know the correct spatial and temporal expression by identifying the enhancers and the transcription factors that regulate this developmental program.

To address this question, in this study we perform transcriptional analysis of *ds* expression. With a combination of experimental approaches, we provide evidence showing that the cis-regulatory elements located in *ds* intronic regions regulate the expression of a diversity of Ds proteins in embryos and larvae, and that some of these isoforms are stage-specific. Furthermore, using the *Drosophila* wing as model to perform a functional analysis, our results suggest that wing growth and PCP are regulated by different Ds proteins expressed at the same time in the wing disc. Furthermore, we can distinguish different regions of the wing disc in which Ds proteins regulate either growth or PCP, and we also predict a differential spatial pattern of the Ds proteins in the developing wing disc.

## RESULTS

### Novel stage-specific Ds isoforms expressed in *Drosophila* embryos

In *Drosophila,* the *ds* gene spans approximately 80kb of genomic DNA on chromosomal region 2L. However, the *ds* locus could be potentially larger since two large deficiencies which uncover adjacent genomic DNA regions such as *Df(2L)al* and *Df(2L)S2* (Flybase), fail to complement *ds* alleles and cause severe phenotypes (Fig. 1A; (Rodriguez, 2004). At present, up to 15 different exons separated by introns that vary over a wide size range (from 0.8 to 48kb) have been identified (Fig. 1B) (Clark et al., 1995). They give rise to four alternative transcriptional variants, DsFL, DsEx, Ds1 and DsIntra, expressed in imaginal discs and brain.

To explore whether the diversity of Ds isoforms expressed in larval tissues also occurred at earlier developmental stages, we analyzed by Western blot whole extracts from wild-type embryos using two specific antibodies raised against epitopes from the cytoplasmic (Ds^cyt^) and the extracellular region (Ds^ex^) of DsFL protein (Fig. S1) (Revilla-Yates et al., 2015);(Strutt and Strutt, 2002). A preliminar analysis showed the detection of multiple bands indicating the expression of several Ds proteins during embryogenesis (Fig. 1C; red frame). Moreover, the protein profiles detected with Ds^cyt^ and Ds^ex^ antibodies differed in the number and size of the bands (Fig. 1C; compare Ewt lanes in both red frames). Three proteins, named E1, E2 and E3, were more abundantly expressed. Indeed, these embryonic isoforms differ in their protein structure: whereas E1 protein was only detected with Ds^cyt^, E3 was detected with Ds^ex^ (Fig. 1C; red asterisk). Furthermore, the isoform E2 (< 50 kDa), detected with both antibodies, would correspond to a small transmembrane protein with both domains, extracellular and cytoplasmic, similar to the large isoform DsFL. Strikingly, E2 and E3 were not found in protein extracts from larval tissues (imaginal discs and brain), suggesting that different Ds proteins are expressed in these two developmental stages (Fig. 1C; compare lanes Ewt and Lwt).

With these evidences it was not possible to discern whether the detected Ds proteins were transcriptional variants or derived from post-translational modifications. Therefore, we analyzed protein extracts from *ds^D36^* embryos and *ds^38K^* larvae, two regulatory alleles that cause embryonic and larval lethality, respectively (Fig. 1C; see Material & Methods). In both mutant conditions, the expression levels of specific protein bands were altered, suggesting that they should correspond to transcriptional isoforms of Ds. Indeed, proteins such as E2 showed a marked decrease in *ds^D36^* embryos (Fig. 1C; compare lanes Ewt and Eds^D36^). Similarly, the levels of the transcriptional isoforms Ds1 and DsIntra, were also reduced in *ds^38K^* larvae (Fig. 1C; compare lanes Lwt and Lds^38k^).

Surprisingly, in *ds^38K^* extracts (Lds^38K^) several bands were overexpressed with respect to wild-type (Lwt), suggesting that this regulatory allele drastically modified the transcription status of *ds* gene allowing the expression of transcripts silenced in wild-type cells (Fig. 1C, blue dot). We have further investigated the transcriptional activity in *ds^38K^* and *ds^D36^* mutants analysing by qRT-PCR the expression of individual exons (Fig. S2). A significant decrease in the expression of most exons was observed compared to control animals. Exceptionally exon2, that encodes part of the extracellular region, showed higher levels indicating an increased transcriptional usage of this exon that correlate with the higher expression of proteins detected with Ds^ex^ antibody (Fig. 1C, blue dot). We did not attempt to determine the sequence of novel isoforms due to experimental constrains to know the full-length cDNA sequence, a crucial step for protein structure prediction. One is the limited length of the cDNA fragments amplified by RACE technique (< 2 kb) to determine unambiguously the 5’ and 3’ ends of the novel complete cDNAs, and the other is the problem of distinguishing among different transcriptional variants sharing common exons when they are coexpressed in the same tissue.

Together, these results reveal the presence of novel Ds isoforms expressed during *Drosophila* embryonic development. The three embryonic proteins expressed at higher levels, E1, E2 and E3, showed differences in their protein structure. At least two of them are not detected in larval tissues, therefore indicating the existence of stage-specific Ds isoforms. Furthermore, the decreased expression observed in *ds^D36^* embryos by Western blot and the qRT-PCR results suggest that at least certain embryonic Ds proteins would correspond to transcriptional isoforms. However, we cannot rule out that undetermined post-translational modifications also contribute to the diversity of the Ds proteins detected in embryos.

**Figure 1.**
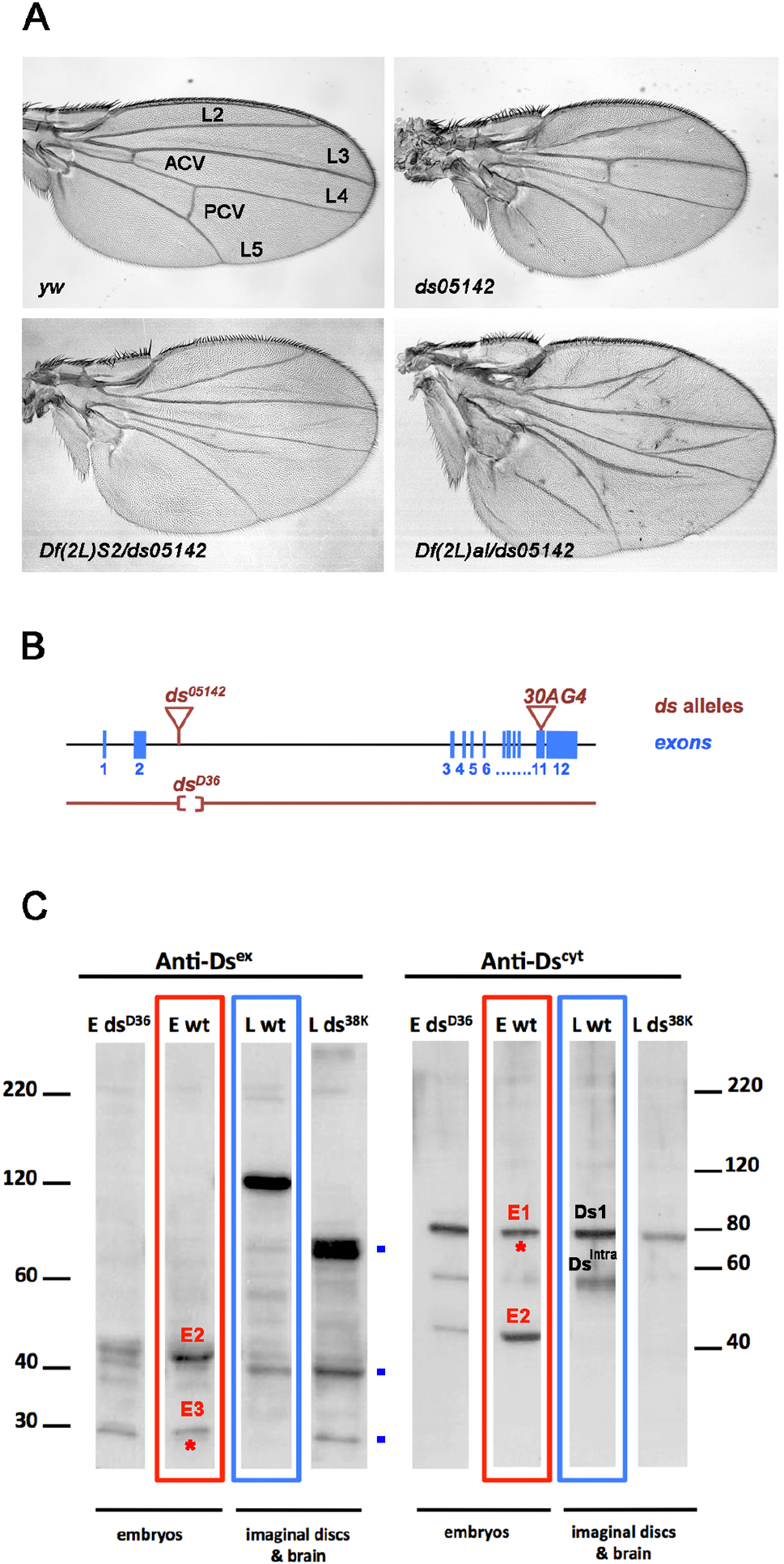
Stage-specific Ds proteins expressed during Drosophila development. (**A**) Representative wings of female flies of yw (control), *ds^05142^* and two heteroallelic combinations of two large deficiencies *Df(2L)S2* and *Df(2L)al* with *ds^05142^* allele. L2 to L5: longitudinal veins. ACV: anterior crossvein. PCV: posterior crossvein. (**B**) Schematic drawing of *ds* gene indicating the relative position of *ds^05142^, ds^D36^* and *30AG4* alleles respect to the exons and intronic regions shown as blue boxes and black lines respectively. (**C**) Western blot analysis of whole extracts from wildtype, *ds^D36^* embryos and *ds^38k^* larvae using two specific antibodies raised against different epitopes of DsFL protein. Ewt and Eds^D36^ lanes correspond to whole extracts derived from wild-type and *ds^D36^* embryos. Lwt and Lds^38K^ lanes correspond to whole extracts from wild-type and *ds^38k^* third instar larvae imaginal discs and brains. Lanes enclosed by a red square correspond to protein profiles of wild-type embryos detected with Ds^ex^ (extracellular region) and Ds^cyt^ (cytoplasmic region) antibodies. Three predominant protein bands are indicated as E1, E2 and E3. Lanes enclosed by a blue square correspond to protein profiles of wild-type larvae detected with Ds^ex^ and Ds^cyt^ antibodies. Ds1 and DsIntra are two Ds proteins previously characterized in larval imaginal discs and brains. Red asterisks mark the embryonic proteins only detected by one of the two Ds antibodies. Blue dots mark the protein bands overexpressed in *ds^38K^* larvae respect to wt larvae (Lwt). Note that these bands were only detected with Ds^ex^ antibody indicating that the corresponding proteins do not contain sequences from the cytoplasmic region.

### Intronic cis-regulatory elements regulate the spatial and temporal expression of *ds* gene

The expression of the *<u>ds</u>* gene is very dynamic and highly localized in imaginal discs and brain (Rodriguez, 2004); (Dearborn and Kunes, 2004). Both *ds* mRNA and protein show a similar expression pattern that is recapitulated by the *lacZ* expression of the *ds^05142^* allele (Fig. 2A; see Materials & Methods). Less known is the expression pattern of *ds* during embryogenesis. Similar to larvae, *ds* mRNA showed a restricted expression in different tissues throughout embryonic development (Fig. 2B). After a strong maternal expression, *ds* mRNA subsequently became more restricted to specific domains along the anterior-posterior (A/P) body axis in segments of the head, thorax, and abdomen. Next, we analyzed *ds^05142^* embryos, where the *lacZ* expression was almost absent at early stages, and only from stage 11 it was detected in certain domains of endogenous *ds* expression. This suggests that certain regulatory elements are located too far to influence this reporter gene during embryonic development (Fig. 2B and Fig. S1). Together, these results showed that *ds* is expressed in multiple cell types and exhibits a dynamic pattern throughout development, suggesting a tight control of its temporal and spatial expression.

To identify regulatory elements of *ds* expression, we started searching in the intronic regions (IR). Several indirect evidences suggest that introns might contain cis-regulatory elements to control its expression in embryos and larvae. First, the *ds^D36^* allele, a small deletion within the large intron 2 (> 48 kb), causes embryonic lethality, the most severe mutant phenotype to date. Second, *ds^05142^* mutant wings exhibit a severe phenotype affecting growth, PCP and patterning, indicating that this P-element insertion disrupt *ds* expression (Fig. 2C; Table S1) (Rodriguez, 2004);(Matakatsu and Blair, 2004). Since the intronic regions of the *ds* gene extend over more than 60 kb of genomic DNA, a strategy of stepwise deletions was not indicated. Instead, we analysed lines from Janelia Collection (Pfeiffer et al., 2008). These lines express GAL4 under the control of genomic DNA fragments derived from intronic regions of the *Drosophila* genome. We have examined fourteen Janelia lines containing intronic sequences of *ds* gene (see Materials & Methods). These lines contain DNA fragments from introns 1, 2, 3, 4, 5 and 10 (Fig. 3A, 3B). Using the *UAS-GFP* reporter line we have illustrated the spatial expression pattern in imaginal discs and brain of third instar larvae of six Janelia Gal4 lines (Fig. 3). Five of them (IR1, IR2A, IR2B, IR3 and IR10) expressed GFP revealing the presence of enhancer elements (Fig. 3C). The IR4 and IR5 lines, that comprise the whole sequence of introns 4 and 5, did not show any obvious GFP expression, indicating that these introns do not contain enhancer activity for these tissues (Fig. 3C). On the contrary, the IR2B and IR10 lines displayed ectopic GFP expression in subdomains where *ds* mRNA is not expressed (Fig. 3C, white arrowheads) suggesting the existence of negative regulators that might contribute to refine the spatial pattern in specific tissues. Strikingly, none of the lines drove GFP expression in regions such as the prospective notum or the eye disc where *ds* mRNA is expressed, indicating that existence of cis-regulatory elements that have not been identified yet (compare Fig. 2A and Fig. 3C).

Together these results show that the intronic regions harbor relevant regulatory elements controlling embryonic and larval *ds* expression. We have mapped some cis-acting regulatory elements in the introns 1, 2, 3 and 10 that direct expression to the brain and imaginal discs. Three of them are tissue-specific enhancers: IR2A and IR10 in imaginal discs and IR2B in the brain. Nonetheless, additional cis-regulatory elements remain unidentified and could be located at long distance from the *ds* promoter, as suggested by the phenotypes observed in allelic combinations with *Df(2L)al* and *Df(2L)S2*.

**Figure 2.**
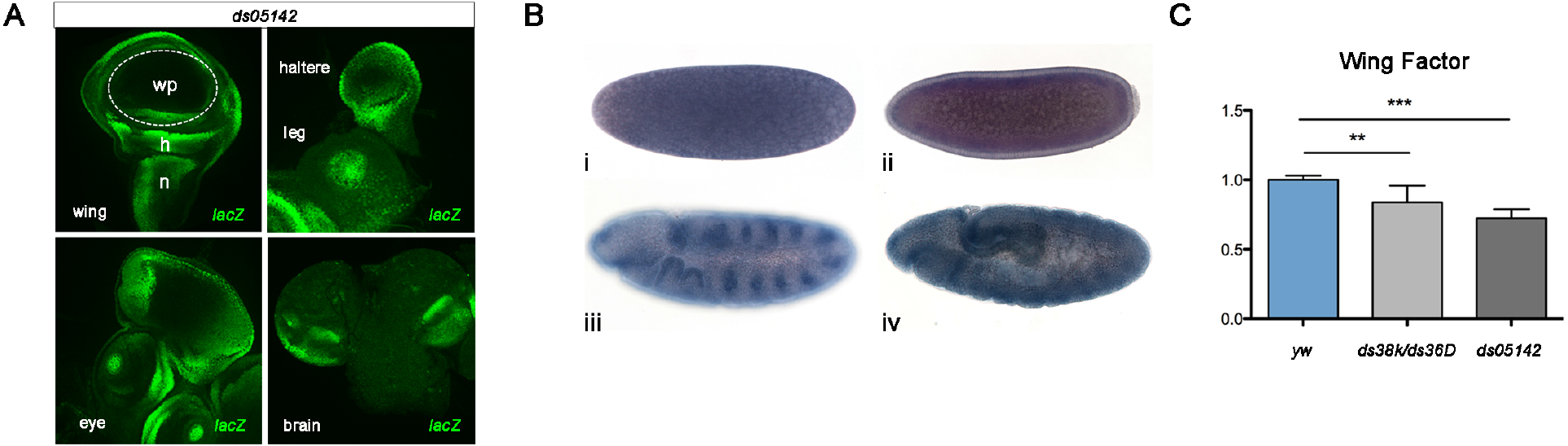
Restricted spatial pattern of *ds* gene at different developmental stages. (**A**) Pattern of *lacZ* expression (green) in the imaginal discs and brain of *ds^05142^* third instar larvae. **wp:** wing pouch; **h:** hinge; **n:** notum. (**B**) Whole-mount *in situ* hybridization of *ds* mRNA in control embryos. Maternal expression of *ds* mRNA was detected at (i) preblastoderm (stage 2). Lateral views of embryos at (ii) cellular blastoderm stage showed a ubiquitous expression. (iii) At stage 11, the expression becomes restricted to a segmented expression pattern in the lateral epidermis and (iv) in endodermal cells that will give rise to the gut tube. Embryos were oriented anterior to the left, dorsal side up. (**C**) Wing area of *ds^05142^, ds^38K^/ds^D36^* and *yw* (control) adult females were quantified using Wing Factor as an arbitrary value (see Table S1). Data were normalized to values obtained with control animals (n ≥ 5 per genotype). Bars depict mean values, and error bars represent S.D. ** p < 0.01, *** p < 0.001.

**Figure 3.**
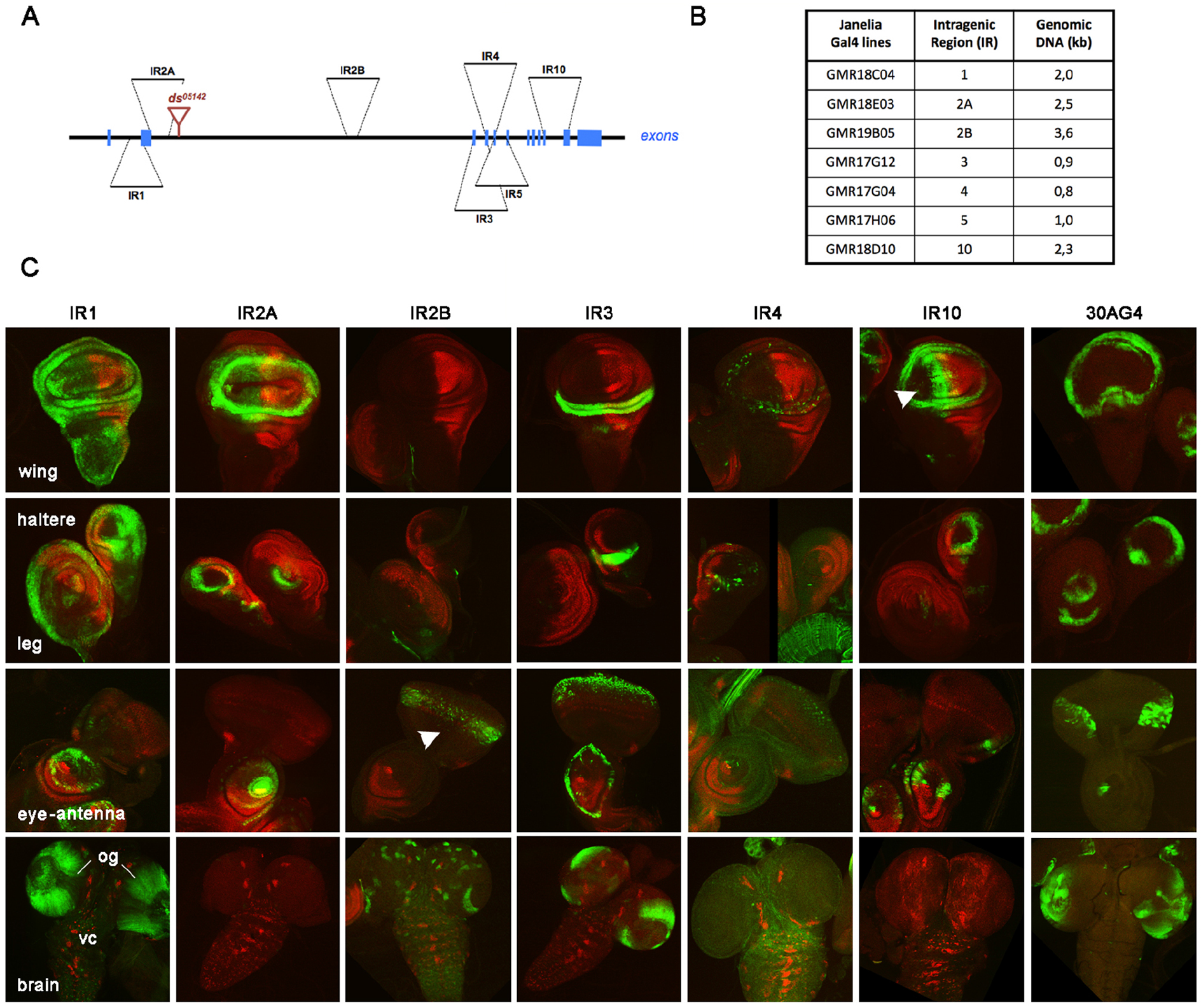
Spatial expression of *ds* is controlled by intronic regulatory elements. (**A**) Schematic representation of the *ds* gene indicating the relative position of the intronic sequences analyzed (IR). (**B**) Summary table showing the sizes of the genomic DNA fragments and the intronic location of Janelia GAL4 lines. (**C**) Expression pattern in the imaginal discs (wing, haltere, leg, eye-antenna) and brains of third-instar larvae. *UAS-GFP* was used as reporter gene (green). In IR5 line, GFP expression was undetectable in imaginal discs and brain (not shown). White arrowheads point to the ectopic GFP expression observed in the eye-antenna disc (morphogenetic furrow) of IR2B line and the wing disc (A cells adjacent to A/P compartment border) of IR10 line. Posterior compartment was visualized with Engrailed (red). **vc:** ventral chord. **og:** optical ganglia.

### The Ds proteins regulating growth and PCP are expressed in different domains of the wing disc

In wild-type wing discs, *ds* is expressed in specific domains of the prospective wing, hinge and notum regions, and the global expression pattern is apparently identical when detected with Ds^ex^ and Ds^cyt^ antibodies (Fig. S1). However, this symmetry is broken in *ds^1^* and *ds^38k^* mutant wing discs, suggesting that different Ds isoforms coexist during wing development (Revilla-Yates et al., 2015). Previous assays based in the overexpression of Ds isoforms such as DsFL, DsEx, and DsIntra have shown a differential role of these proteins in the control of wing growth and PCP/patterning (Revilla-Yates et al., 2015). However, nothing is known about the spatial and temporal requirements for the Ds isoforms expressed during wing formation.

To address this point, we analysed, in adult wings, the contribution of Ds proteins to the control of PCP and growth by depleting *ds* expression in different domains of the wing disc (Fig. 4).

For this purpose, we expressed two independent, isoform-specific, RNAi lines under the control of endogenous regulatory elements such as IR1, IR2A, IR3 or *30AG4,* an enhancer-trap line inserted in exon11, and two additional Gal4 lines: *salPEG4* and *nubG4* expressed in broader domains of the prospective wing territory (Fig. 4). For each genotype, the hair polarity (Fig. 4A’-F’’) and the wing area were measured (Fig. 4G-L; Table S1). The dsRNAi-ex and dsRNAi-cyt lines targeted sequences encoding for the extracellular and cytoplasmic regions of Ds, respectively. Their efficiency to silence *ds* expression had been previously tested by quantitative RT-PCR (Revilla-Yates et al., 2015).

We found that wings of flies expressing dsRNAi-cyt in the central wing pouch driven by Gal4 lines from different sources (IR1, IR2A, *nubG4* and *salPEG4*) exhibited hair misorientation and visible patterning defects, both phenotypes associated with disruption of PCP signalling (Fig. 4A’,4B’,4E’,4F’). In contrast, the expression of dsRNAi-ex in the same conditions resulted in apparently normal wings (Fig. 4A’’,4B’’,4E’’,4F’’) (Revilla-Yates et al., 2015). Remarkably, although *ds* is expressed at high levels in the proximal wing and hinge (Fig. 2A), neither dsRNAi-cyt nor dsRNAi-ex caused detectable alterations in PCP when they were expressed in these domains (IR3 and *30AG4*) (Fig. 4C’,4D’,4C’’,4D’’). These results indicate that the PCP/patterning function is exerted by those Ds proteins expressed in the prospective wing pouch.

Next, in order to determine the effects in wing growth, we measured the surface of wings expressing dsRNAi-cyt and dsRNAi-ex (Fig. 4G-L). In the proximal wing and hinge (IR3 and *30AG4),* the expression of dsRNAi-ex and dsRNAi-cyt caused a significant reduction of the wing size suggesting that the Ds proteins expressed in the periphery of the wing pouch are involved in the positive regulation of wing growth without affecting PCP (Fig. 4I, 4J).

By contrast, in the wing pouch territory, depending on which RNAi line was expressed, they have opposite effects on wing size (*nubG4* and *salPEG4*). Whereas flies expressing dsRNAi-ex had wings smaller than controls, similar to what was observed at the periphery of the wing pouch, the depletion of Ds proteins containing a cytoplasmic domain (dsRNAi-cyt) gave rise to enlarged wings (Fig. 4K, 4L). These unexpected results suggest that, in the prospective wing pouch, several Ds isoforms are expressed that contribute to the control of wing size.

Several conclusions can be drawn from these results; one is the existence of several Ds proteins expressed in the developing wing disc that can control either PCP or wing growth. In addition, these proteins have different structural features (extracellular and cytoplasmic domains) and spatial patterns. Second, our results identify the wing pouch as the region in which *ds* exerts the control on PCP and patterning. And third, the expression of different Ds protein isoforms can regulate growth with opposed effects on wing size.

**Figure 4.**
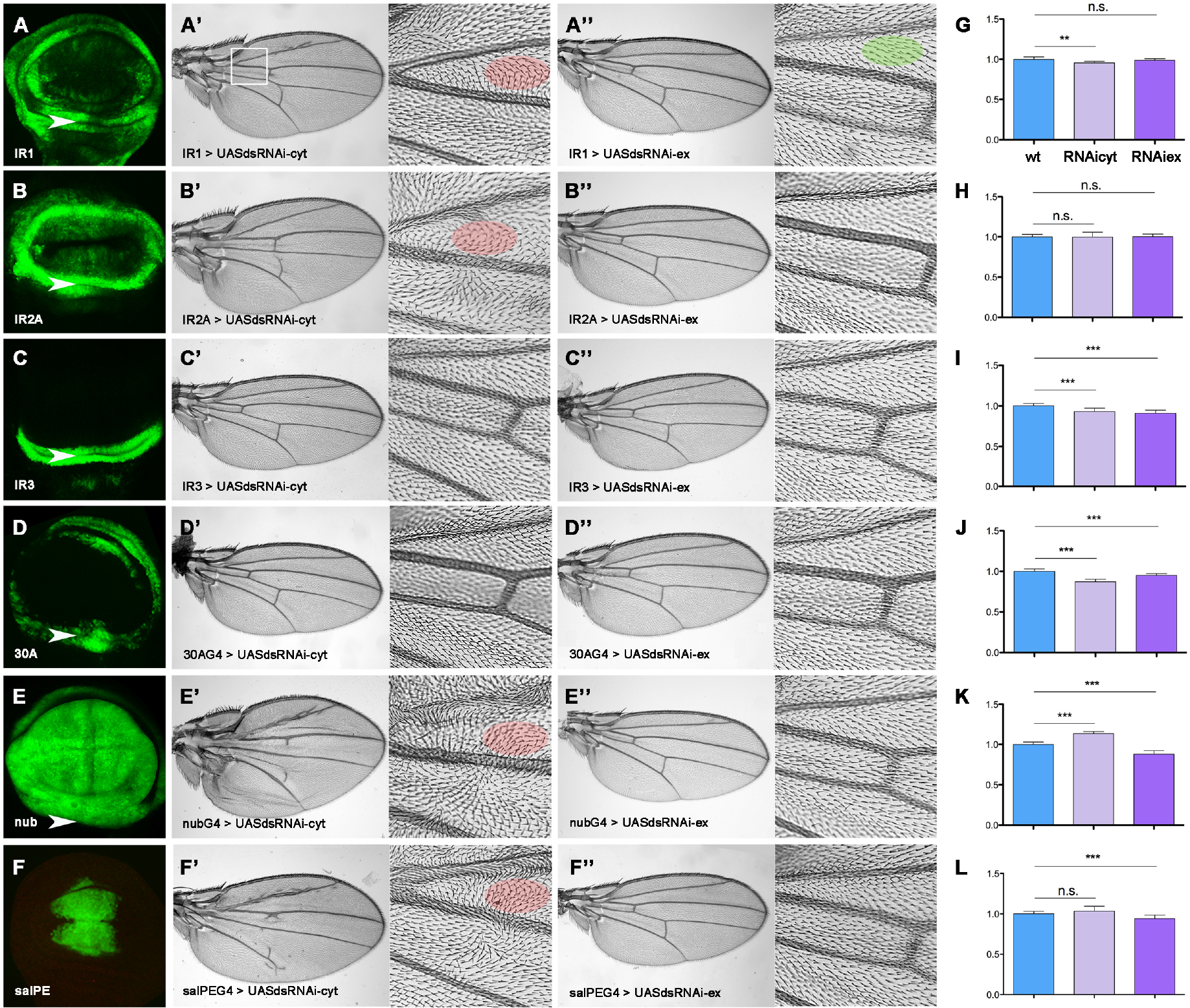
Different Ds proteins regulate Growth and PCP in the wing disc. (A-F) GFP expression pattern of Gal4 lines IR1, IR2A, IR3, *30AG4, nubG4* and *salPEG4* used as drivers to express *UAS-dsRNAi-cyt* and *UAS-dsRNAi-ex.* The white arrowheads indicate the border between the proximal wing and the hinge territory. Representative images of wings expressing (A’-F’) UAS-dsRNAi-cyt and (A’’-F’’) UAS-dsRNAi-ex in different regions of the wing imaginal disc. Close-up image of the proximal region (white square) of adult wings is shown for each genotype to visualize the hair swirling pattern associated to PCP defects. Hair (trichome) misorientation was highlighted by a red oval. The green oval showed the normal hair orientation pointing towards to the distal end of the wing. Only flies expressing *dsRNAi-cyt* in the central wing pouch (IR1, IR2A, *nubG4* and *salPEG4*) exhibit PCP phenotypes such as hair misorientation and visible patterning defects. Expression of dsRNAi-ex and dsRNAi-cyt in the proximal wing and hinge (IR3 and *30AG4*) caused a significant reduction in wing size. However, *dsRNAi-cyt* expression in the prospective wing pouch (*nubG4* and *salPEG4*) has an opposed effect in growth giving rise to enlarged wings. (G-L) Graphs show the quantification of the wing area of female flies for each genotype (n ≥ 5). Bars depict mean values, and error bars represent S.D. ** p < 0.01, *** p < 0.001, n.s: not significant.

## DISCUSSION

Tissue morphogenesis is a crucial biological process that will result in functional organs with characteristic shapes by the assembly of different cell types into cellular sheets. This process is governed in time and space during development by key regulators. One of these regulators is *ds,* an evolutionarily conserved gene that plays a key role in tissue development and tumorigenesis by controlling different cellular functions, such as the orientation of cell division, cell-cell communication, and cellular differentiation, proliferation, migration and apoptosis.

Several studies have shown the role of *ds* in organ formation by regulating the activity of different signalling pathways in the control of PCP, tissue growth and patterning, in which different Ds proteins are involved (Matakatsu and Blair, 2006); (Willecke et al., 2008); (Baena-Lopez et al., 2008);(Degoutin et al., 2013);(Revilla-Yates et al., 2015). Together, these data showed a scenario much more complex than the one initially thought for the *ds* gene. To better understand how diverse Ds proteins expressed in one tissue can control diverse cellular processes through different molecular mechanisms, here we have focused on the transcriptional regulation of the *ds* gene. Our present analysis shows a variety of Ds protein isoforms expressed at different developmental stages. In embryos, we detected at least two small proteins henceforth named DsE2 and DsE3, that are stage-specific Ds isoforms since they are not expressed in larval tissues. Moreover, the correlation between the protein and mRNA levels (lower or higher) observed in two regulatory alleles such as *ds^D36^* and *ds^38K^*, further support that some of these embryonic and larval Ds isoforms are transcriptional variants. Second, we have shown the relevance of the intronic regions for the regulation of *ds* expression in embryos and larval tissues. We have unveiled the activity of intronic cis-regulatory elements in the introns 1, 2, 3 and 10 that control the dynamic and restricted expression pattern of *ds* in larval brain and imaginal discs and rule out enhancer activity for these tissues in introns 4 and 5. In contrast, although the embryonic lethality associated to *ds^D36^* allele support the presence of regulatory elements in intron2 to control *ds* expression at this developmental stage, our results discard the presence of enhancer elements involved in the early *ds* expression pattern in the intronic sequences (IR) analyzed. Therefore, additional enhancer elements still unidentified must be required to fully explain the expression pattern of *ds* at embryonic and larval stages. The *Df(2L)al* and *Df(2L)S2* phenotypes would suggest that some of these regulatory elements might be located in the intragenic regions, at a long distance from the *ds* promoter.

Third, using the *Drosophila* wing as model to perform a functional analysis of Ds proteins, we found that the isoforms expressed in the developing wing differentially contribute to the control of growth and PCP/patterning. The phenotypic analysis of wings in which *ds* expression was depleted by two isoform-specific RNAi lines under the control of endogenous regulatory elements revealed that the cellular functions regulated of by Ds take place in specific domains of the wing disc. Remarkably, the structures of the Ds isoforms involved in each function seem to be different. Our results highlighted the importance of the wing pouch domain for the regulation of wing patterning and PCP. Indeed, from the Ds proteins expressing this domain, those containing the cytoplasmic region are particularly relevant to be in charge of the control of these functions.

In contrast, we show that wing size is controlled by different Ds proteins expressed in both the proximal wing/hinge and the prospective wing pouch. However, they exert opposed effects on wing size since, while those expressed in the wing/hinge positively control wing growth, the proteins expressed in the prospective wing pouch seem to repress cell proliferation. We predict that this specific regional function of Ds proteins largely reflects their restricted expression patterns, although we cannot discriminate individual patterns because the molecular tools available recognize sequences shared in common for more than one isoform. We propose that the participation of different Ds proteins with opposed effects on wing size might be undertaken combining both mechanisms (the cell-autonomous and the non-cell autonomous) to provide a balance between cell proliferation and apoptosis, resulting from the regulated activity of the Hippo pathway and other signalling pathways including JNK and those activated by the morphogens Dpp, Hh and Wg.

Together, these findings suggest that the genomic organisation of the *ds* gene is highly complex in order to ensure the expression of specific Ds isoforms with restricted spatial and temporal patterns at different stages to activate different cell-specific molecular programs through development. Moreover, the present results provide an increased knowledge about the molecular regulation of *ds* gene that will contribute to understanding how *DCHS1* and *DCHS2* genes can control developmental processes in health and disease in humans.

## MATERIALS AND METHODS

### *Drosophila* stocks

The following stocks were obtained from the Bloomington Stock Center (BDSC): *yw^1118^, UAS-GFP, ds^38k^, ds^05142^, Df(2l)al, Df(2L)S2, 30AG4, nubG4* and the Janelia GAL4 lines. *salPEG4* line drives expression specifically to the central domain of the prospective wing pouch (Cruz et al., 2009). *Df(2l)al* (21C1;21C7) and *Df(2L)S2* (21C8;22B1) are large deficiencies that uncover chromosomal regions adjacent to the *ds* locus (21E2). The *ds^38k^* allele (http://flybase.org/reports/FBal0028156.html) has been described as a spontaneous mutation. We have checked by sequencing that the splicing sites and the exons 1 to 12 were intact. *ds^05142^* is a P-lacZ transposable element inserted in intron 2 at 4 kb from the 3’ end of exon1. *ds^D36^* is a small deletion of 0,4 kb in intron 2 that behaves as an embryonic lethal allele (Rodriguez, 2004). *30AG4* is a GAL4 enhancer-trap line inserted in exon 11 of *ds* gene and behaves as a weak hypomorphic allele (Revilla-Yates et al., 2015).

The Janelia lines express the Gal4 protein under the control of genomic DNA fragments derived from non-coding regions of the *Drosophila* genome (Pfeiffer et al., 2008). These constructs were inserted in the same orientation at a known genomic location by site-specific integration to have the same genomic position effects in the pattern of GAL4 expression for all the lines. In this study we used as Gal4 drivers the following lines mapping within *ds* locus: <u>GMR18C04,</u> GMR18E03, GMR19B05, <u>GMR17G12,</u> <u>GMR17G04</u>, <u>GMR17H06</u>, and GMR18D10 described in Flybase (https://bdsc.indiana.edu/stocks/gal4/gal4janelia.html). The underlined stocks are currently not available at BDSC but the information is found in the Enhancer sequence section of BDSC.

The RNAi lines from the Vienna Drosophila RNAi Center (VDRC), UAS-dsRNAi-cyt (14350) and UAS-dsRNAi-ex (2646) specifically target sequences (exons 12 and 6) encoding for the extracellular and cytoplasmic domains respectively. The efficiency of both RNAi lines has been tested by qRT-PCR assays (Revilla-Yates et al., 2015). Potential off-targets were not identified by *in silico* prediction. Flies were maintained at 25°C on standard *Drosophila* medium. For experiments with embryos, females were allowed to lay eggs for 24 h on apple juice agar plates with yeast. *yw^118^* was used as control in all experiments.

### Adult wing analysis

For the analysis of the wing area, female adult wings were dissected and mounted in a solution of lactic acid:ethanol (1:3). Images of the wings were acquired with a Zeiss Axiovert200 microscope. The surface area and the distance between the distal ends of L2 and L5 veins were traced using Adobe Photoshop CS3 for the following genotypes: *yw^1118^, ds^38K^/ds^D36^, ds^05142^* and the *UAS-dsRNAi-cyt* and *UAS-dsRNAi-ex* strains crossed to the GAL4 lines GMR18C04, GMR18E03, GMR17G12, 30AG4, nubG4, salPEG4. To calculate the wing area of *yw^1118^, ds^38K^/ds^D36^, ds^05142^* wings we used the wing factor. Wing factor is an arbitrary value calculated as the ration between the wing surface area (size) and the distance between the distal ends of L2-L5 veins (shape). For the remaining genotypes, the wing area was calculated directly from the value obtained using Adobe Photoshop and normalized respect to control wings. The mean value of the wing area for each genotype is shown in Supplementary Information, Table S1.

### In situ hybridization and immunohistochemistry

Whole-mount in situ hybridisation with digoxigenin-labelled DB1 RNA probe and immunohistochemistry were performed as described previously in (San Martin and Bate, 2001). *Drosophila yw* and *ds^05142^* embryos were collected at 25°C. The RNA probe was prepared according to the manufacturers instructions (Roche). DB1 is a fragment of *ds* cDNA described in (Clark et al., 1995). Imaginal discs and brain were dissected, fixed and stained as described (Gomez-Skarmeta et al., 1995). Primary antibodies used were: rabbit anti-β-galactosidase (1:10000; Cappel), mouse anti-Engrailed (1:50 Developmental Studies Hybridoma Bank, DSHB), rabbit anti-Ds (Ds^ex^ 1:1000 a gift from D.Strutt (Strutt and Strutt, 2002) and guinea pig anti-Ds (Ds^cyt^ 1:500 (Revilla-Yates et al., 2015). Images were acquired with a Zeiss Axiophot microscope and confocal microscope LSM710 (Zeiss). Images were processed using Adobe Photoshop CS4 and ImageJ software.

### Quantitative real-time PCR analysis

Total RNA from three independent biological samples of each genotype and developmental stage was extracted using Trizol (Invitrogen). The first cDNA strand was synthesized with the Transcriptor First Strand cDNA Synthesis kit (Roche) using 2 μg of total RNA and oligo (dT) primer. To quantify the expression of individual exons all PCR reactions were carried out in technical duplicates using HotStart Taq polymerase (Qiagen), and SYBR green (Qiagen). Data were acquired using a 7900HT Fast Real-Time PCR System (Applied BioSystems). Reactions were normalized to the expression level of *Elf-2a* and *Tbp* genes used as endogenous controls. The primer sequences and amplicon size for individual exons are described in Revilla-Yates et al. 2015. Exon12 was split into two fragments 12A and 12B. Fragment 12B comprises the DNA sequences corresponding to the cytoplasmic region. For relative quantification, the normalization of biological samples was performed using the comparative Ct (2-ΔΔCt) method after efficiency calculation for each primer pair. Data analysis was carried out using Integromics RealTime StatMiner software (http://www.integromics.com/StatMiner).

### Statistical analysis

In all the experiments pertaining to wing area and qRT-PCR, differences in the mean value were assessed using the unpaired two-tailed Student’s t-test. Boxes represent the mean value. *p* value of <0.05 was considered statistically significant. Data are presented as relative to the mean of *yw^1118^* used as control.

### Western blot analysis

Protein extracts were obtained from embryos and from imaginal discs and brain of third instar larvae. Tissue was dissected in Ringer´s solution, supplemented with a complete protease inhibitor cocktail (Roche). Protein extracts were separated on a 10% Mini-PROTEAN TGX Precast Gel (BioRad) and transferred onto PVDF membranes (BioRad). For detection, the primary antibodies used were guinea pig anti-Ds^cyt^ (1:2000) and rabbit anti-Ds^ex^ (1:2000). Antirabbit and anti-mouse HRP-conjugated secondary antibodies were used 1:2000. Membranes were developed with ECL reagent kit.

## ACKNOWLEDGEMENTS

We thank Juan Modolell, Sonsoles Campuzano and Joaquim Culi for critical reading of the manuscript and valuable discussion. We also thank M. Gómez for her advice and help with the qRT-PCR experiments and data analysis. We acknowledge Helen and David Strutt for Ds^ex^ antibody. *Drosophila* fly stocks were obtained from the Bloomington Drosophila Stock Center (NIH P40OD018537) and VDRC Stock Center and the antibodies from Developmental Studies Hybridoma Bank. This work was funded by a programme grant from the Spanish Ministry of Economy (BFU2008-01869) to IR, a Fellowship to ERY from Word Works Science and an institutional grant from the Fundación Ramon Areces to the CBMSO.

## REFERENCES

Baena-Lopez, L. A., Rodriguez, I. and Baonza, A. (2008). The tumor suppressor genes dachsous and fat modulate different signalling pathways by regulating dally and dally-like. Proc Natl Acad Sci U S A 105, 9645–50.

Bando, T., Mito, T., Maeda, Y., Nakamura, T., Ito, F., Watanabe, T., Ohuchi, H. and Noji, S. (2009). Regulation of leg size and shape by the Dachsous/Fat signalling pathway during regeneration. Development 136, 2235–45.

Cappello, S., Gray, M. J., Badouel, C., Lange, S., Einsiedler, M., Srour, M., Chitayat, D., Hamdan, F. F., Jenkins, Z. A., Morgan, T. et al. (2013). Mutations in genes encoding the cadherin receptor-ligand pair DCHS1 and FAT4 disrupt cerebral cortical development. Nat Genet 45, 1300–8.

Chung, S., Vining, M. S., Bradley, P. L., Chan, C. C., Wharton, K. A., Jr. and Andrew, D. J. (2009). Serrano (sano) functions with the planar cell polarity genes to control tracheal tube length. PLoS Genet 5, e1000746.

Clark, H. F., Brentrup, D., Schneitz, K., Bieber, A., Goodman, C. and Noll, M. (1995). Dachsous encodes a member of the cadherin superfamily that controls imaginal disc morphogenesis in Drosophila. Genes Dev 9, 1530–42.

Clemenceau, A., Berube, J. C., Belanger, P., Gaudreault, N., Lamontagne, M., Toubal, O., Clavel, M. A., Capoulade, R., Mathieu, P., Pibarot, P. et al. (2017). Deleterious variants in DCHS1 are prevalent in sporadic cases of mitral valve prolapse. Mol Genet Genomic Med 6, 114–120.

Cruz, C., Glavic, A., Casado, M. and de Celis, J. F. (2009). A gain-of-function screen identifying genes required for growth and pattern formation of the Drosophila melanogaster wing. Genetics 183, 1005–26.

Dearborn, R., Jr. and Kunes, S. (2004). An axon scaffold induced by retinal axons directs glia to destinations in the Drosophila optic lobe. Development 131, 2291–303.

Degoutin, J. L., Milton, C. C., Yu, E., Tipping, M., Bosveld, F., Yang, L., Bellaiche, Y., Veraksa, A. and Harvey, K. F. (2013). Riquiqui and minibrain are regulators of the hippo pathway downstream of Dachsous. Nat Cell Biol 15, 1176–85.

Fabre, C. C. G., Casal, J. and Lawrence, P. A. (2008). The abdomen of Drosophila: does planar cell polarity orient the neurons of mechanosensory bristles? Neural Dev 3, 12.

Gomez-Skarmeta, J. L., Rodriguez, I., Martinez, C., Culi, J., Ferres-Marco, D., Beamonte, D. and Modolell, J. (1995). Cis-regulation of achaete and scute: shared enhancer-like elements drive their coexpression in proneural clusters of the imaginal discs. Genes Dev 9, 1869–82.

Matakatsu, H. and Blair, S. S. (2004). Interactions between Fat and Dachsous and the regulation of planar cell polarity in the Drosophila wing. Development 131, 3785–94.

Matakatsu, H. and Blair, S. S. (2006). Separating the adhesive and signaling functions of the Fat and Dachsous protocadherins. Development 133, 2315–24.

Matis, M. and Axelrod, J. D. (2013). Regulation of PCP by the Fat signaling pathway. Genes Dev 27, 2207–20.

Pfeiffer, B. D., Jenett, A., Hammonds, A. S., Ngo, T. T., Misra, S., Murphy, C., Scully, A., Carlson, J. W., Wan, K. H., Laverty, T. R. et al. (2008). Tools for neuroanatomy and neurogenetics in Drosophila. Proc Natl Acad Sci U S A 105, 9715–20.

Rawls, A. S., Guinto, J. B. and Wolff, T. (2002). The cadherins fat and dachsous regulate dorsal/ventral signaling in the Drosophila eye. Curr Biol 12, 1021–6.

Revilla-Yates, E., Varas, L., Sierra, J. and Rodriguez, I. (2015). Transcriptional analysis of the dachsous gene uncovers novel isoforms expressed during development in Drosophila. FEBS Lett 589, 3595–603.

Rodriguez, I. (2004). The dachsous gene, a member of the cadherin family, is required for Wg-dependent pattern formatión in the Drosophila wing disc. Development 131, 3195–3206.

San Martin, B. and Bate, M. (2001). Hindgut visceral mesoderm requires an ectodermal template for normal development in Drosophila. Development 128, 233–42.

Sotos, J., Miller, K., Corsmeier, D., Tokar, N., Kelly, B., Nadella, V., Zhong, H., Wetzel, A., Adler, B., Yu, C. Y. et al. (2017). A patient with van Maldergem syndrome with endocrine abnormalities, hypogonadotropic hypogonadism, and breast aplasia/hypoplasia. Int J Pediatr Endocrinol 2017, 12.

Strutt, H. and Strutt, D. (2002a). Nonautonomous planar polarity patterning in Drosophila: dishevelled-independent functions of frizzled. Dev Cell 3, 851–63.

Willecke, M., Hamaratoglu, F., Sansores-Garcia, L., Tao, C. and Halder, G. (2008). Boundaries of Dachsous Cadherin activity modulate the Hippo signaling pathway to induce cell proliferation. Proc Natl Acad Sci U S A 105, 14897–902.

Zwaveling-Soonawala, N., Alders, M., Jongejan, A., Kovacic, L., Duijkers, F. A., Maas, S. M., Fliers, E., van Trotsenburg, A. S. P. and Hennekam, R. C. (2018). Clues for Polygenic Inheritance of Pituitary Stalk Interruption Syndrome From Exome Sequencing in 20 Patients. J Clin Endocrinol Metab 103, 415–428.

